# CardioWAS Pathway-Based RNA-Seq and Genome Integration Platform for CVD Omics-Wide Associations

**DOI:** 10.1101/2025.10.06.679779

**Authors:** Yuvraj Singh, Nikhil Sharma, Kabir Raghuvanshi, Jennifer Kwan, Alokkumar Jha

## Abstract

The integration of transcriptomic and genomic data is essential for dissecting the molecular basis of complex cardiovascular diseases (CVD). Existing resources primarily focus on differential gene expression at the individual gene level, with limited support for pathway-level interpretation or genomic integration. We introduce CARDIOWAS (Cardiovascular Omics-Wide Association Analysis Platform), the first Shiny-based, cardiovascular-specific platform that performs pathway-centric expression profiling in conjunction with Genome-Wide Association Study (GWAS) integration. CARDIOWAS systematically aggregates and harmonizes RNA-seq data from 18 NCBI GEO studies, comprising 419 heart failure (HF) and 208 control samples, and applies robust batch correction followed by DESeq2-based differential expression analysis. Gene-level statistics are translated into pathway effect sizes using curated KEGG and Reactome annotations, enabling systematic evaluation of disease-relevant biological modules.

CARDIOWAS extends beyond transcriptomic profiling by incorporating GWAS-identified loci, mapping variants to proximal genes and pathways, and leveraging machine learning models to highlight transcriptome–variant–pathway associations. Both germline and somatic variant contributions can be interrogated, providing a framework to link non-coding variation with altered pathway activity. Application of the platform identified 12 HF-specific and 19 CAD-enriched pathways, implicating inflammatory, metabolic, and signaling processes central to disease biology. Implemented as a user-friendly Shiny interface, CARDIOWAS allows researchers to upload transcriptomic data, visualize perturbed pathways, evaluate variant burden, and prioritize druggable targets. By uniting large-scale RNA-seq data with GWAS evidence in a pathway-centric framework, CARDIOWAS represents a novel, reproducible, and accessible tool for precision medicine in CVD and offers a resource for omics-driven hypothesis generation and translational research.

## Introduction

Cardiovascular diseases (CVD), including heart failure (HF) and coronary artery disease (CAD), remain the leading causes of morbidity and mortality worldwide^1^. Despite significant advances in clinical diagnostics and therapeutics, our understanding of the molecular and genetic basis of CVD progression and treatment response remains limited^2,3^. A major challenge in this field is the ability to integrate transcriptomic and genomic data at a systems level to uncover disease-driving pathways and potential therapeutic targets. Traditional differential expression analyses focus on identifying gene-level changes between disease and control conditions. However, these approaches often fail to capture the broader context of molecular interactions and do not account for the regulatory and pathway-level perturbations that underlie complex diseases. Moreover, the disconnection between transcriptome data and Genome-Wide Association Study (GWAS) findings limits our ability to interpret the functional consequences of non-coding or regulatory variants, especially those residing in somatic regions. To bridge this gap, we introduce CARDIOWAS (Omics-Wide Association Analysis Platform) — a comprehensive platform designed to integrate Transcriptome-Wide Association Studies (TWAS)^4^ with GWAS findings to facilitate pathway-specific analyses in cardiovascular research. Unlike existing tools, CARDIOWAS goes beyond simple differential gene expression by focusing on the aggregation of gene-level effects into biologically meaningful pathways^5^. It further enhances interpretability by integrating somatic and germline variant data from the GWAS Catalog^6^, using machine learning to uncover variant-transcript-pathway relationships. CARDIOWAS aggregates publicly available datasets from NCBI GEO and harmonizes data across 18 independent studies involving HF and CAD. By applying rigorous normalization, batch correction, and statistical modeling, CARDIOWAS provides a robust framework to identify differentially expressed genes (DEGs), compute pathway-level effect sizes, and correlate these with disease-associated genetic variants. Importantly, CARDIOWAS supports user-friendly, patient-level analyses through a Shiny-based web application, empowering researchers to conduct translational investigations with minimal bioinformatics expertise. Our platform represents a critical step forward in systems-level cardiovascular genomics. By integrating transcriptomic changes with genetic variation at the pathway level, CARDIOWAS enables researchers to move beyond isolated biomarkers and toward a more holistic understanding of disease biology, paving the way for personalized risk assessment and targeted therapeutic strategies.

## Methods

### 2.1 Data Collection and Preprocessing

Publicly available bulk RNA-seq datasets were identified in the NCBI Gene Expression Omnibus (GEO)^7^ using search terms related to heart failure (HF), cardiomyopathy, and coronary artery disease (CAD) (Table 1). Studies were retained if they included clearly labeled HF and control samples, provided raw count or count-compatible processed matrices, and contained sufficient metadata for harmonization. Eighteen independent studies met inclusion criteria, yielding 419 HF and 208 control samples. Gene identifiers were standardized to current HGNC symbols; duplicated mappings were resolved by collapsing counts only when gene model equivalence was verified. Samples were excluded if metadata were incomplete, if more than 30% of genes had zero counts, or if quality control (QC) metrics indicated outlier status based on library size, proportion of lowly expressed genes, and Cook’s distance from a preliminary negative-binomial fit. A unified count matrix was constructed, and batch effects were addressed with ComBat-seq^8^ using study accession as the batch factor and disease status as a protected variable. Size-factor normalization used the median-of-ratios method. Two working representations were generated: normalized counts for hypothesis testing and variance-stabilized transformed (VST) values for visualization and machine learning. QC included principal component analysis and hierarchical clustering before and after batch correction to verify attenuation of batch-driven structure. Genes expressed in fewer than 10% of samples at counts per million greater than one was removed to reduce sparsity.

### 2.2 Differential Expression and Pathway Scoring

Differential expression was performed with DESeq2^9^ using a design formula including batch and condition (HF versus control). Dispersion estimates were fit to a parametric trend, and Wald tests were used for contrasts. P-values were adjusted using the Benjamini–Hochberg procedure. Genes were considered differentially expressed at false discovery rate (FDR) < 0.05 with absolute log2 fold-change > 1. For cross-sample comparability, gene-wise Z-scores were computed from VST values within study and re-centered on the merged cohort.

Pathway mapping used curated KEGG^10^ and Reactome^11^ gene sets. For each pathway, a signed activity score was computed as the mean Z-score across member genes. Uncertainty was estimated by nonparametric bootstrap across samples with five thousand resamples, and empirical P-values were derived from size-matched random gene sets that preserved gene-wise variance characteristics. Resulting P-values were FDR-adjusted across all tested pathways. Pathway scores were standardized to zero mean and unit variance for downstream modeling.

### 2.3 Integration with GWAS and Feature Construction

Variant–trait associations were retrieved from the GWAS Catalog and filtered for cardiovascular traits, including HF, CAD, lipids, blood pressure, and inflammation biomarkers. Variants were mapped to genes using a hybrid strategy combining nearest transcription start site proximity (±100 kb), overlap with promoter and enhancer annotations, and reported eQTL targets when available. Variant–gene links were joined to transcriptomic statistics and propagated to pathways via the curated gene sets. Somatic mutation regions were defined by intersecting variant coordinates with catalogs of recurrent somatic hotspots and clonal hematopoiesis loci from public resources; a binary indicator captured whether a GWAS variant or a high-linkage disequilibrium proxy overlapped these regions. The feature space for modeling comprised pathway activity scores, selected gene-level Z-scores, variant burden indicators at gene and pathway levels, somatic-region flags, study indicators to capture residual structure, and available clinical covariates (e.g., age and sex) encoded and centered. Continuous features were scaled on training folds only, with parameters applied to held-out data.

### 2.4 Machine Learning Modeling and Validation

Classification of HF versus control status used a random forest algorithm within a nested, reproducible pipeline. To mitigate information leakage, feature selection was performed exclusively within training folds based on FDR and effect size thresholds, and highly collinear features (absolute Pearson correlation > 0.9) were pruned. Model development used stratified five-fold cross-validation repeated ten times. Hyperparameters—number of trees, maximum depth, number of variables tried at each split, and minimum node size—were tuned via grid search within each repetition. Class imbalance was handled using inverse-prevalence sample weights, with synthetic minority oversampling applied to training data when class ratio exceeded 1.5:1.

Performance metrics included area under the receiver operating characteristic curve, area under the precision–recall curve, accuracy, sensitivity, specificity, F1-score, and calibration slope. Ninety-five percent confidence intervals were obtained from cross-validation replicates and stratified bootstrap on out-of-fold predictions. Feature importance was assessed by permutation, and marginal effects for top features were summarized using partial-dependence and accumulated local effects profiles. To test robustness against residual batch structure, leave-one-study-out validation was conducted by iteratively holding out each study as an independent test set.

### 2.5 Statistical Software and Reproducibility

All analyses were executed in R (version ≥ 4.3). Core packages included DESeq2 for differential expression, sva for ComBat-seq, data.table and dplyr for data processing, and ranger or randomForest for modeling. ROC and PR analyses were conducted with pROC and PRROC^12^, and model interpretation utilized iml. Gene set handling used curated GMT import utilities and msigdbr where appropriate. Workflow orchestration used targets or drake for dependency-tracked execution with deterministic random seeds. Package versions were pinned via renv, and session information was archived at each pipeline run. All data splits, preprocessing parameters, and tuned hyperparameters persisted to disk for exact re-execution.

### 2.6 Shiny Application for Interactive, Reproducible Exploration

An interactive Shiny application was developed to expose the analysis in a read-only, parameterized interface suitable for exploratory use by collaborators. The application followed a modular architecture comprising (i) a data layer, (ii) an analysis layer, and (iii) a presentation layer. Precomputed artifacts—including normalized counts, VST matrices, differential expression results, pathway activity matrices, cross-validation predictions, feature-importance tables, and model configuration manifests—were drug-target correlation modules, making CARDIOWAS a continuously evolving platform for translational cardiovascular research.

### Dynamic Transcriptomic Profiling Through Single Dataset Analysis (Uncovering gene-level and pathway-level insights in individual cardiovascular disease datasets.)

To facilitate individualized exploration of transcriptomic data, CARDIOWAS offers a Single Dataset Analysis module that enables users to analyze expression profiles from any NCBI GEO dataset using a simple interface. Users can input GEO accession codes and define experimental groups (e.g., control vs. treated) to initiate differential expression analysis. Using dataset GSE106382 as an example, the module guides users through selecting relevant samples and performing normalization and statistical testing using the limma framework. Figure 2a demonstrates the functionality of this module, where sample metadata—including accession numbers, cell types, and condition labels—is displayed in a structured table for easy selection. Following sample input, users can navigate through multiple analysis tabs, including differential expression gene (DEG) identification, pathway enrichment, and gene ontology categorization. This mode supports detailed exploration of disease-relevant expression signatures, empowering researchers to examine changes at both the gene and pathway level for cardiovascular conditions such as heart failure and coronary artery disease. In addition to basic DEG identification, CARDIOWAS provides functional interpretation of gene expression results through integrated Gene Ontology (GO) annotations^13^. As shown in **Figure 2c**, users can explore the biological functions, molecular activities, and cellular processes associated with each significantly altered gene. For instance, the platform displays the gene symbol, title, and corresponding GO terms related to DNA binding, transcription regulation, signal transduction, and cytoskeletal organization. This functionality enables rapid prioritization of genes involved in disease-relevant pathways.

**Figure 1:**
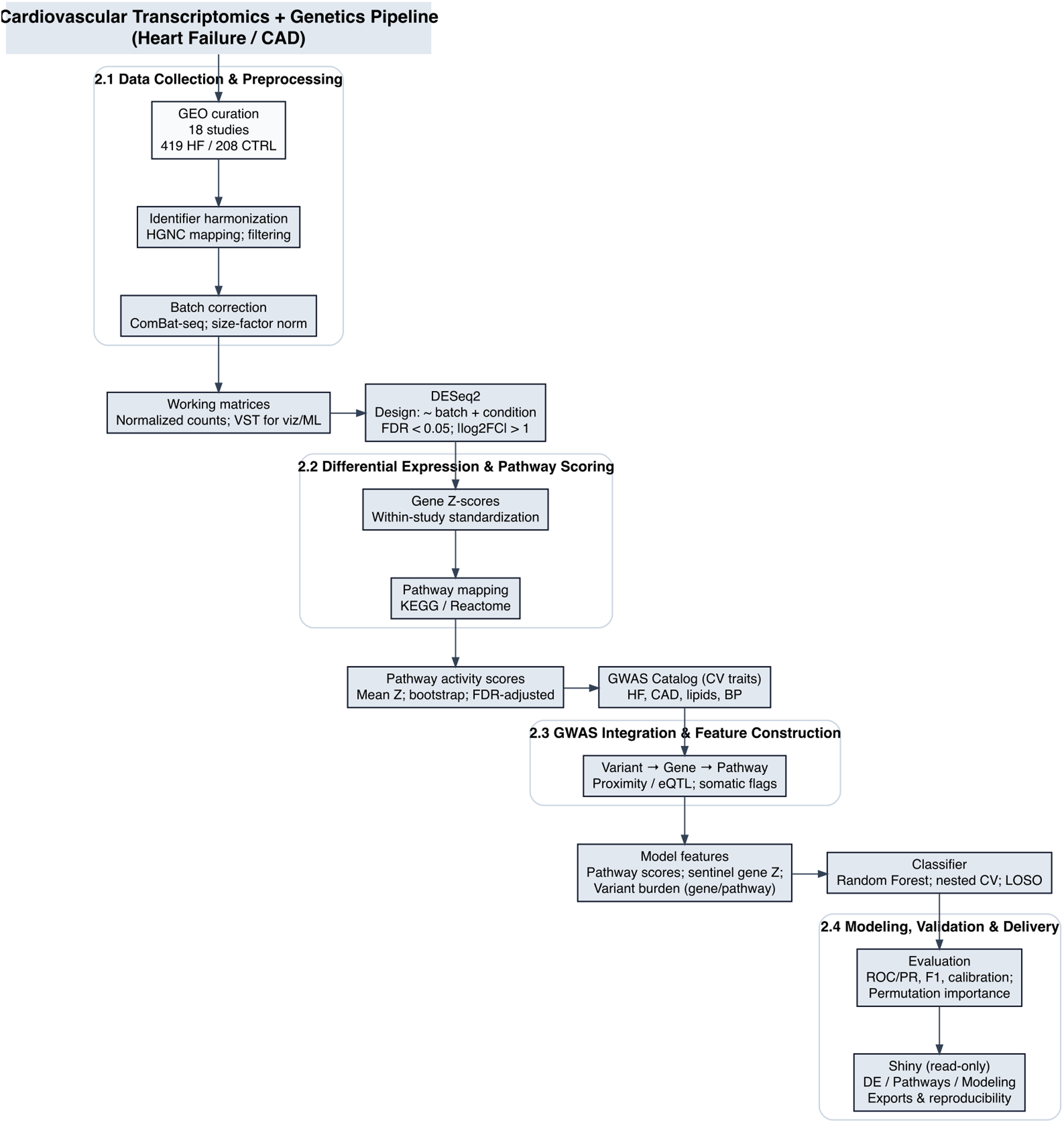
Flow Diagram of Study

**Figure 2.**
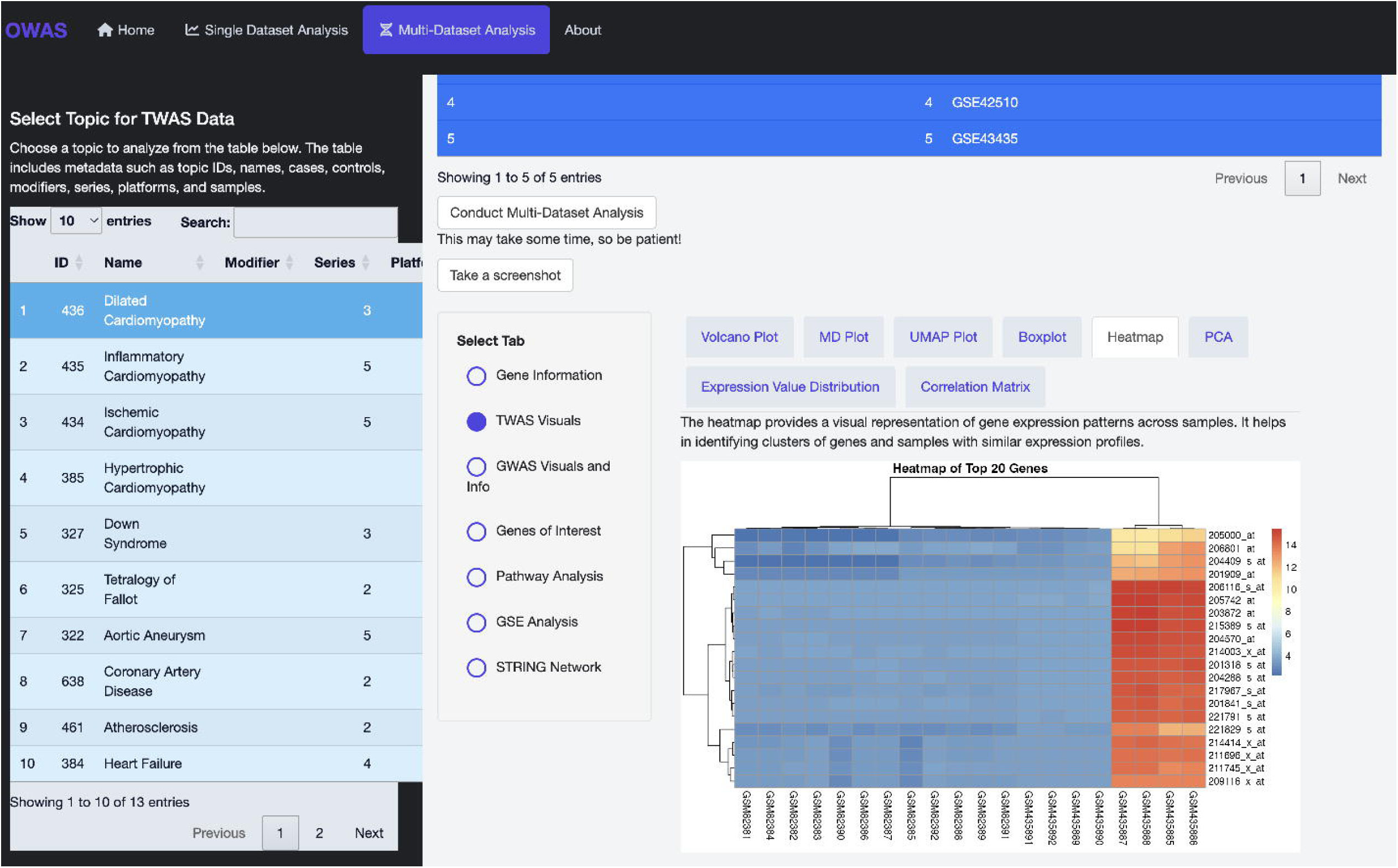
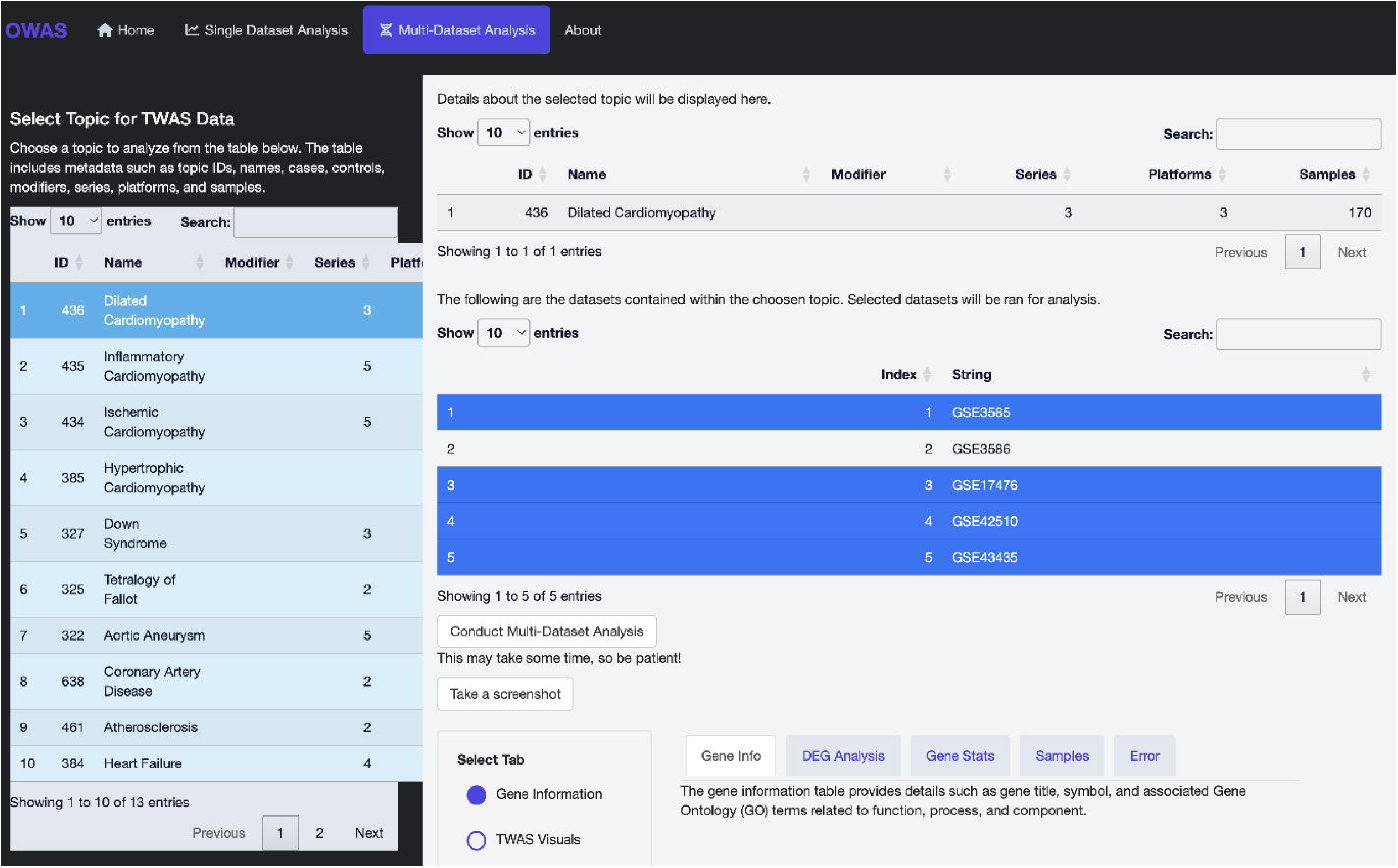
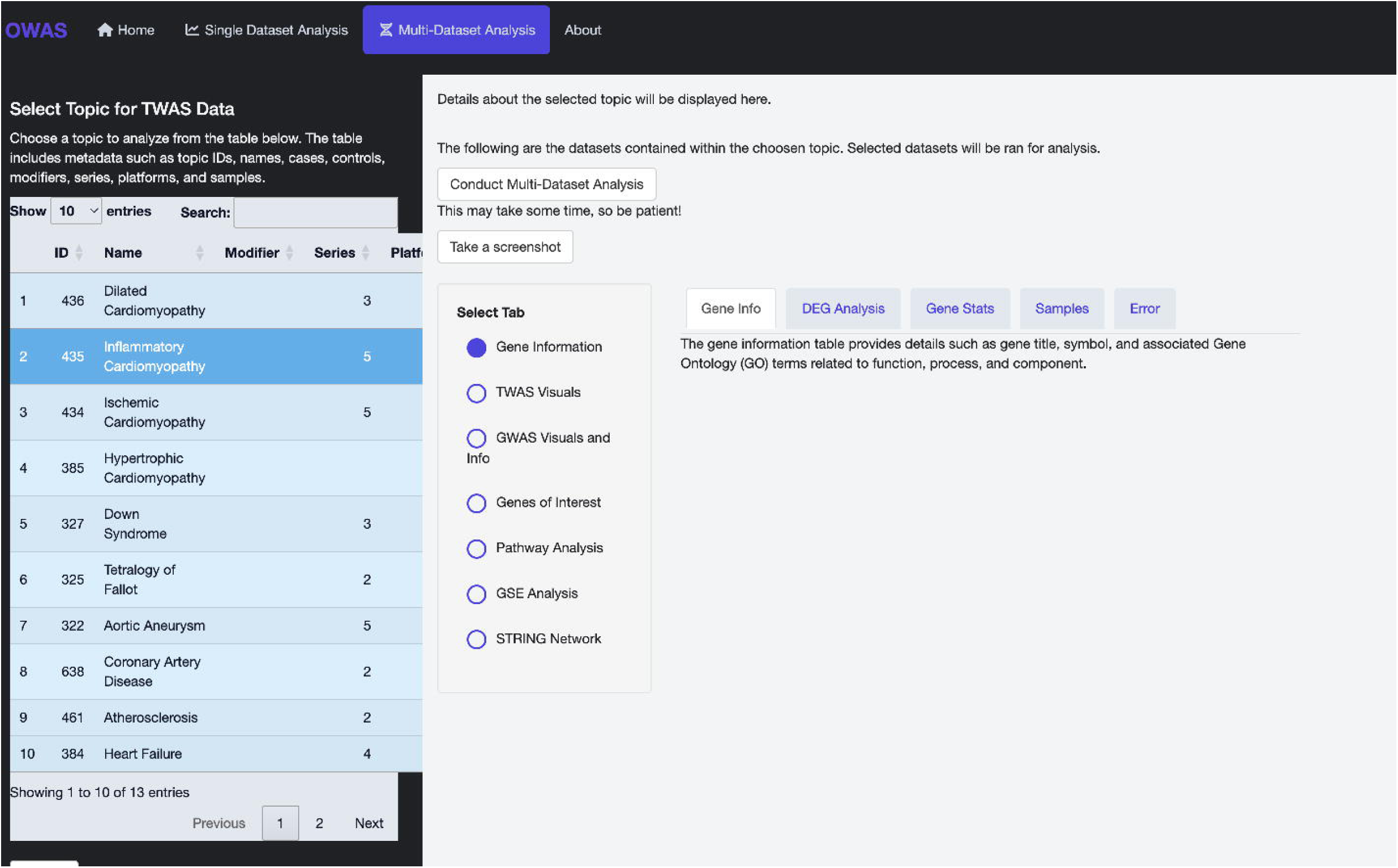
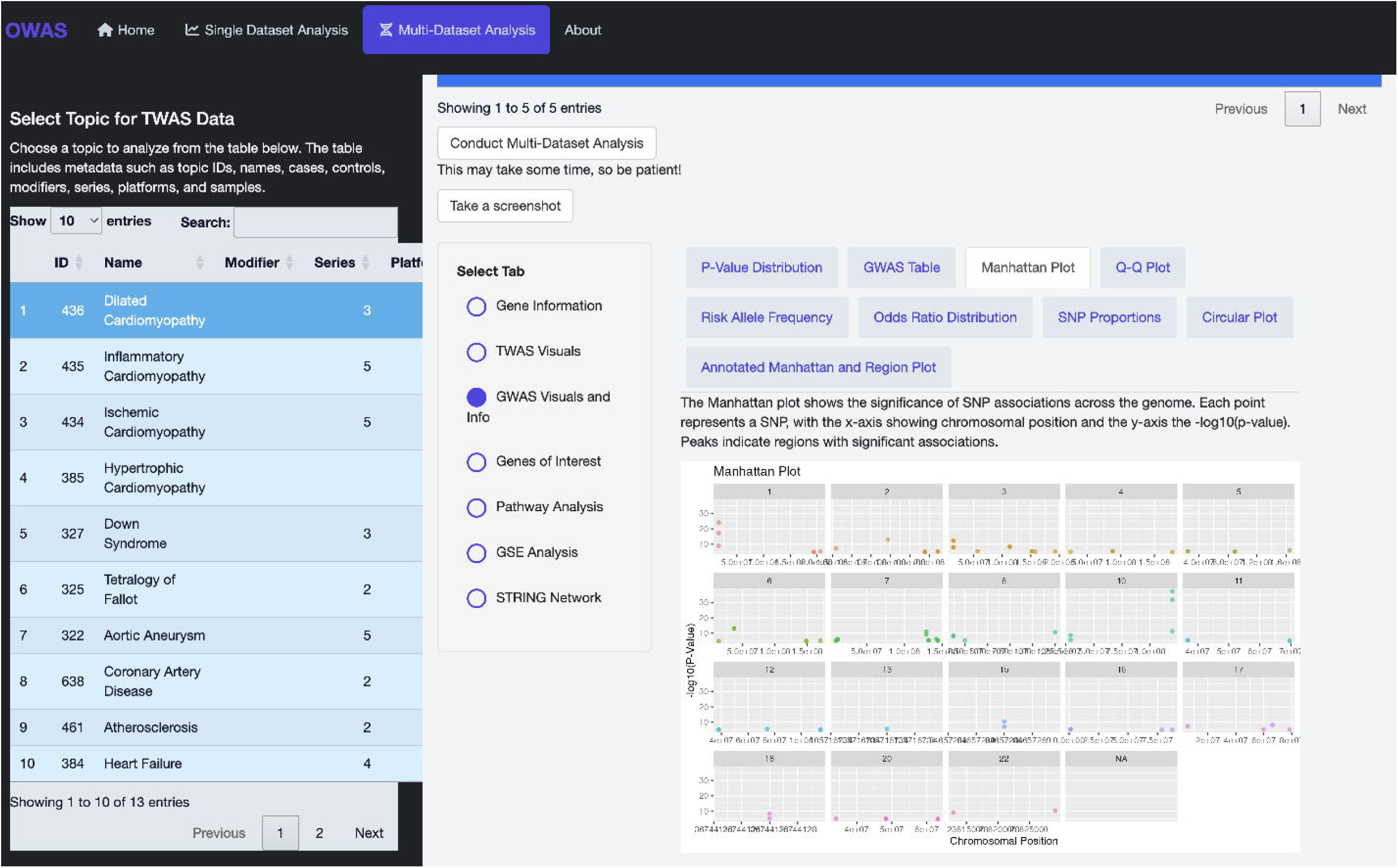
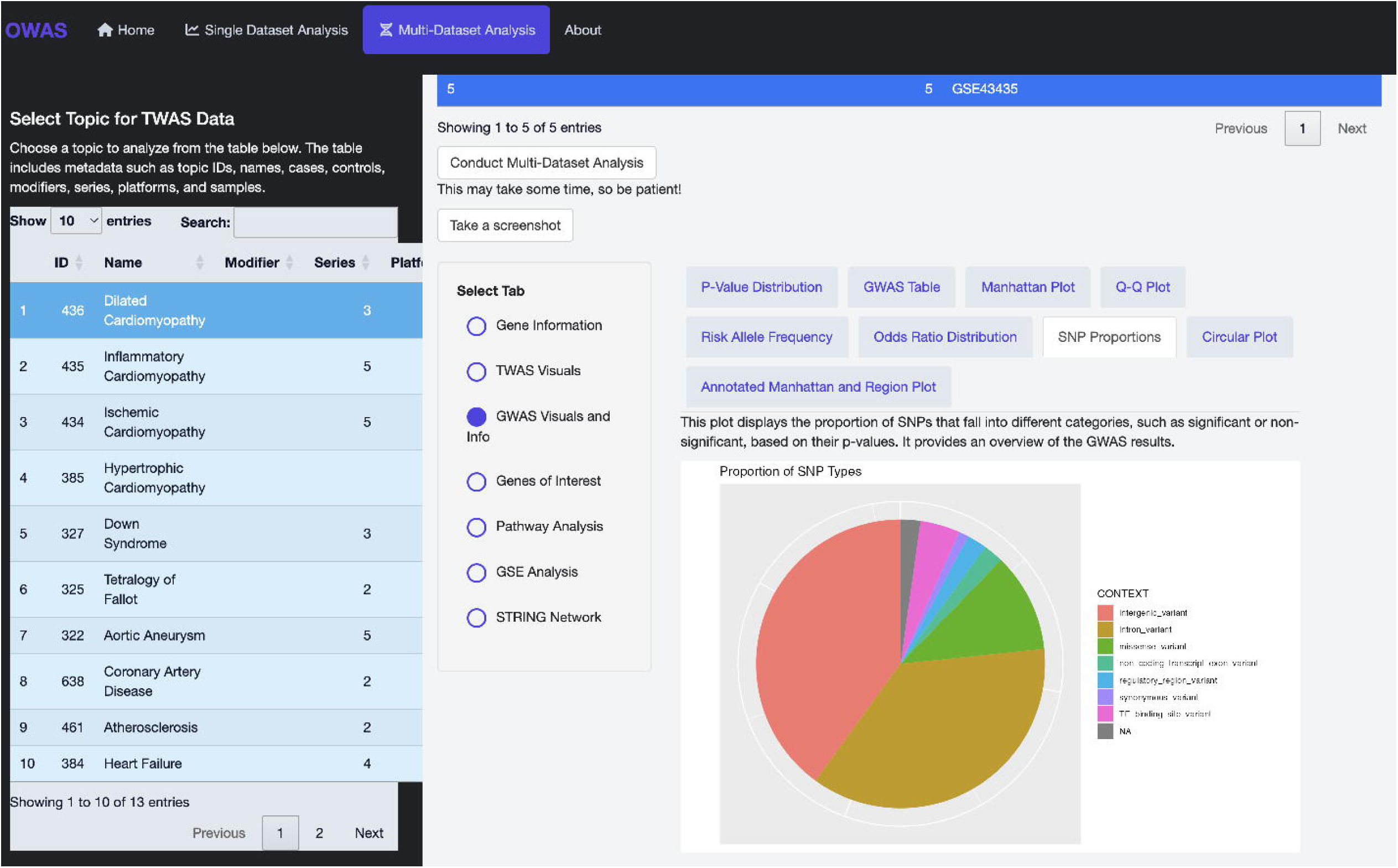
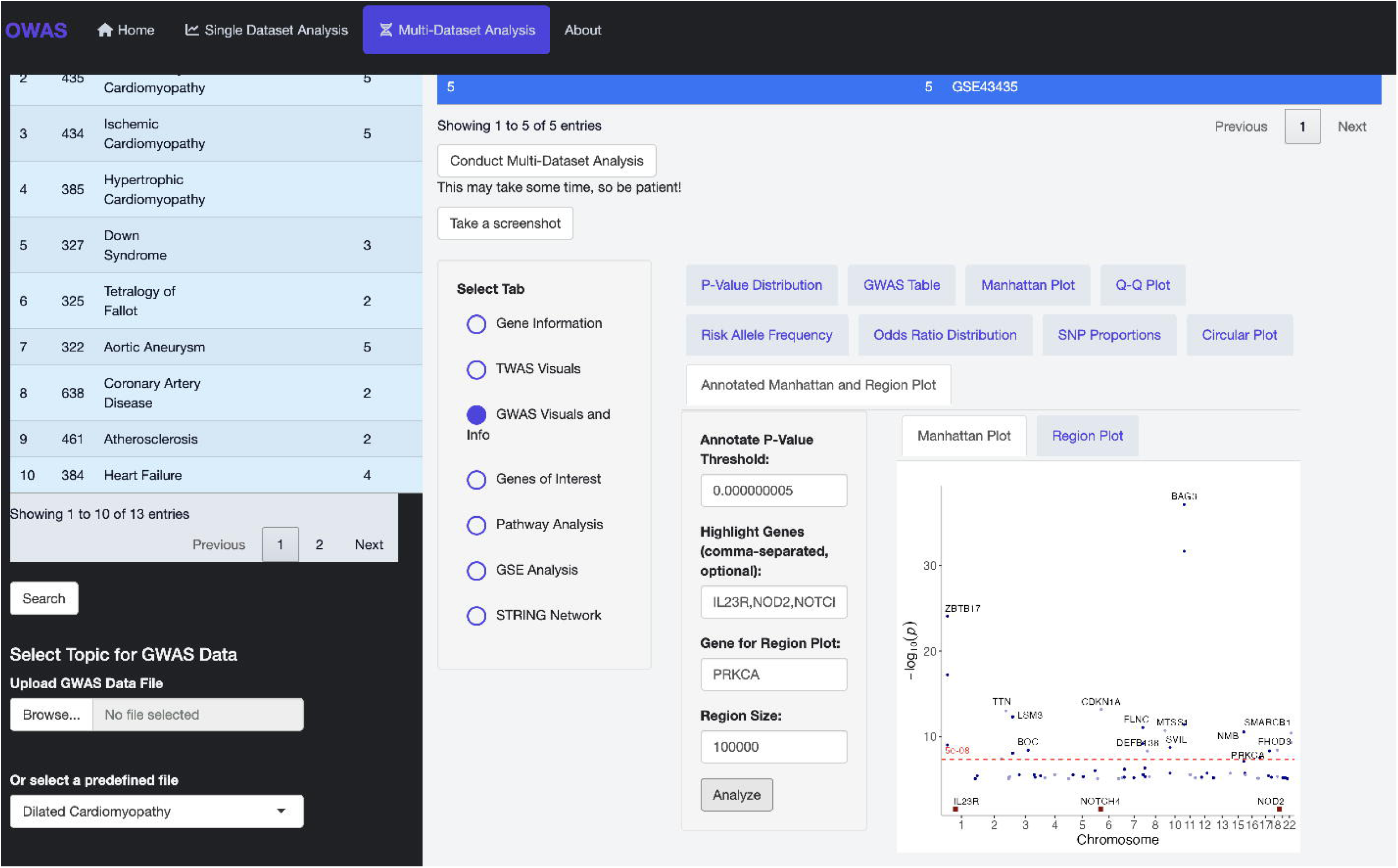
Overview of single dataset analysis functionality in CARDIOWAS using dataset GSE106382. **(a)** The main interface for the Single Dataset Analysis module, where users input GEO accession codes, define experimental groups (e.g., Control vs. Treated), and select relevant samples based on metadata. **(b)** The Gene Information tab showing differentially expressed genes along with their gene symbols, GO molecular functions, and biological processes, providing immediate biological context for DEGs. **(c)** Volcano plot displaying the log2 fold change versus -log10 adjusted p-values for all genes. Genes with high statistical significance and large expression changes are easily identified for prioritization. **(d)** Correlation matrix showing pairwise gene expression similarity across samples. High intra-group correlation suggests consistent biological signals and sample quality. **(e)** Pathway enrichment analysis of differentially expressed genes using the ENRICHR API and KEGG pathways. Several immune, vascular, and metabolic pathways are enriched in the disease group. **(f)** STRING-based protein-protein interaction network of filtered DEGs, highlighting potential gene clusters and hubs such as ENTPD1 and KLRD1 that may drive disease-relevant biological processes.

The gene table can be filtered or exported in various formats (CSV, Excel, PDF), supporting downstream analyses and integration with external tools. The intuitive layout helps users navigate complex functional annotations with ease, allowing cardiovascular researchers to rapidly link altered genes with processes relevant to cardiac remodeling, inflammation, or vascular integrity. To assist with the interpretation of expression data, CARDIOWAS offers a robust set of visualization tools within the *Gene Visuals* module^14^. Among these, the volcano plot provides an intuitive representation of the statistical significance and magnitude of differential gene expression. As illustrated in Figure 2c, each point corresponds to an individual gene, plotted according to its log2 fold change (x-axis) and -log10 adjusted p-value (y-axis). This allows users to easily identify genes that are both significantly and substantially up- or downregulated between experimental conditions.

In the example dataset (GSE106382), clear patterns of differential expression are observed, with a subset of genes displaying high statistical significance and large effect sizes. These visual cues are critical for prioritizing genes for further analysis, such as pathway enrichment or variant annotation. In combination with additional plots (e.g., PCA, heatmap, UMAP), the volcano plot helps users explore dataset structure, variance, and disease-relevant transcriptional shifts in cardiovascular research.

CARDIOWAS also provides tools to assess sample relationships and overall expression patterns using correlation matrices and dimensionality reduction plots. As shown in **Figure 2d**, the correlation matrix visualizes the pairwise correlation coefficients between samples in dataset GSE106382. Each cell represents the degree of similarity between two samples based on global gene expression profiles, with values ranging from −1 (perfect negative correlation) to +1 (perfect positive correlation). This feature is particularly useful for identifying outlier samples, validating group separations (e.g., Control vs. Treated), and ensuring consistency across replicates. In the example shown, strong intra-group correlation can be observed among the majority of samples, suggesting stable transcriptional responses within each condition. Such insights are critical for ensuring robust statistical modeling and improving the biological interpretability of downstream DEG and pathway analyses. To provide deeper biological insights into gene expression changes, CARDIOWAS includes a pathway analysis module that performs enrichment on differentially expressed genes using the ENRICHR API^15^ and curated pathway databases. As shown in Figure 2e, this module displays a bar plot of significantly enriched pathways across KEGG categories. In the example analysis of dataset GSE106382, several pathways emerged as significantly perturbed, including VEGF signaling, vascular smooth muscle contraction, T cell receptor signaling, and Th17 cell differentiation—all relevant to cardiovascular and immune processes. The enrichment results allow users to explore both metabolic and signaling pathways affected in disease states. Each bar in the visualization corresponds to an enrichment score, and selecting a pathway reveals associated gene members and functional annotations. This integrated approach enables researchers to move beyond gene-level observations to biologically meaningful pathway interpretations, facilitating translational discoveries in cardiovascular genomics. Finally, CARDIOWAS provides an integrated STRING network visualization to explore the protein-protein interaction landscape of differentially expressed genes. As demonstrated in Figure 2f, users can set statistical thresholds (e.g., p-value and log fold change cutoffs) to filter the DEG list and generate a gene interaction map. The resulting network displays biological relationships and co-regulation patterns among the most dysregulated genes. In this case, genes such as ENTPD1^16^, KLRD1^17^, and FCYR2^18^ appear as central nodes in the network, indicating potential regulatory hubs relevant to immune signaling and cardiovascular inflammation. This network-based perspective enables researchers to uncover functional gene modules and prioritize candidates for validation or therapeutic targeting. Combined with pathway enrichment and visualization tools, the STRING module provides a systems-level view of gene expression changes, reinforcing CARDIOWAS as a comprehensive tool for cardiovascular omics interpretation.

### Integrated Multi-Dataset Analysis for Cross-Cohort Discovery (*Meta-analysis of cardiovascular transcriptomes to reveal robust, shared pathways and variant associations)*

To enable robust cross-cohort comparison and meta-analysis, CARDIOWAS includes a Multi-Dataset Analysis module, designed for integrating TWAS data across multiple studies and cardiovascular conditions. This module allows users to select from a curated list of disease topics—such as inflammatory cardiomyopathy, heart failure, and aortic aneurysm—each aggregated from multiple datasets with standardized metadata. **Figure 3a** showcases the user interface for selecting and initiating multi-dataset analysis. Researchers can explore a list of conditions and select a phenotype of interest (e.g., Inflammatory Cardiomyopathy), after which the platform automatically retrieves and harmonizes all relevant GEO datasets associated with that condition. Upon initiating the analysis, CARDIOWAS performs batch effect correction, DEG identification, and pathway-level aggregation across studies. The integrated tab system supports detailed examination of TWAS visuals, GWAS overlays, shared genes of interest, and pathway enrichment across cohorts. This functionality empowers users to identify consistent molecular signatures across datasets and disease subtypes, offering a more generalizable view of disease mechanisms than single-study analyses can provide. As illustrated in **Figure 3b**, once a disease topic is selected, CARDIOWAS displays all constituent datasets associated with that phenotype. For example, selecting *Dilated Cardiomyopathy* reveals five independent GEO series (e.g., GSE3585, GSE17476, GSE42510), comprising a total of 170 samples collected across three different platforms. This layout allows users to verify study inclusion and metadata consistency prior to executing batch-integrated analysis. With a single click, the system harmonizes these datasets, correcting for batch effects and platform differences using standardized preprocessing pipelines. This functionality is essential for maximizing statistical power and enhancing reproducibility when studying complex traits across heterogeneous datasets. By combining multiple datasets into a unified analysis pipeline, CARDIOWAS enables cross-validation of findings and facilitates identification of universally dysregulated pathways and biomarkers in cardiovascular disease. Upon completion of multi-dataset analysis, CARDIOWAS generates comprehensive visual summaries to reveal conserved gene expression signatures across datasets. As shown in **Figure 3c**, a heatmap of the top 20 differentially expressed genes highlights the consistency of transcriptomic patterns across samples from five GEO series related to dilated cardiomyopathy. Each row represents a gene, and each column corresponds to a sample, with the color intensity indicating the level of expression. This visualization enables users to identify clusters of co-expressed genes and segregate samples based on shared expression profiles, providing an intuitive overview of transcriptional dysregulation. In this analysis, several genes showed markedly elevated or suppressed expression across datasets, reinforcing their potential as cross-cohort biomarkers. Hierarchical clustering further aids in discovering underlying structure among samples, such as subtypes or progression stages, which might otherwise be masked in single-study analyses. To further strengthen disease gene discovery, CARDIOWAS enables integration of transcriptomic signatures with genome-wide association study (GWAS) data. As illustrated in **Figure 3d**, the *GWAS Visuals and Info* module generates a Manhattan plot summarizing the statistical significance of SNP-trait associations across the genome. Each point represents a single nucleotide polymorphism (SNP), with chromosomal location on the x-axis and –log10(p-value) on the y-axis. Peaks above the significance threshold indicate loci that may harbor disease-relevant genetic variation. In the case of dilated cardiomyopathy, the integrated GWAS output helps prioritize transcriptionally altered genes that also lie near significant SNPs. This enables the identification of potential regulatory variants and enhances confidence in the biological relevance of observed transcriptomic shifts. Additional visualizations such as Q-Q plots, odds ratio distributions, and risk allele frequencies provide further statistical context, creating a holistic view of gene-variant-pathway relationships. To complement statistical significance plots, CARDIOWAS provides categorical summaries of SNP annotations, offering users insight into the biological context of variants. **Figure 3e** presents a pie chart displaying the proportion of SNP types identified in the GWAS data for dilated cardiomyopathy. Each slice represents a variant class—such as intronic, intergenic, missense, synonymous, and regulatory region variants—highlighting the genomic distribution and potential functional impact of the SNPs. This breakdown aids researchers in distinguishing between non-coding variants likely to affect gene regulation and coding variants that may alter protein structure or function. For instance, a high proportion of intronic and regulatory region variants, as seen here, may suggest transcriptional or splicing regulation mechanisms underlying disease phenotypes. By integrating SNP type distributions with expression and pathway results, CARDIOWAS supports comprehensive multi-layered interpretation of omics data for cardiovascular conditions. To further refine the interpretation of GWAS findings, CARDIOWAS includes an annotated Manhattan and region plot module, enabling users to focus on specific genes or genomic regions of interest. **Figure 3f** illustrates the use of this feature, where genes such as *IL23R*, *NOD2*, and *NOTCH4* are highlighted, and *PRKCA* is specified for a regional zoom-in. The resulting Manhattan plot overlays user-defined gene names with corresponding SNP significance values, while the region plot zooms into the genomic neighborhood surrounding *PRKCA*, displaying detailed association signals. This functionality empowers researchers to link GWAS peaks directly to biologically plausible candidate genes and examine the surrounding linkage disequilibrium structure. It is particularly valuable for interpreting non-coding variants and prioritizing genes for functional validation. Through this integrated, interactive visual analytics framework, CARDIOWAS bridges the gap between transcriptome signals, genetic variants, and disease pathways in cardiovascular research. To enhance biological relevance and prioritize functional targets, CARDIOWAS provides a *Genes of Interest* module to identify genes shared between TWAS differential expression and GWAS association results. As shown in **Figure 3g**, users can define statistical thresholds (e.g., adjusted p-value and log fold change) and generate a table of overlapping genes. In the case of dilated cardiomyopathy, two key genes—CCND1 and FLNC—were identified as common to both transcriptomic and genomic signals. These overlapping genes represent high-confidence candidates for further validation, as they are supported by both regulatory and genetic evidence. The module supports export in multiple formats (CSV, Excel, PDF), facilitating downstream pathway mapping or drug repurposing efforts. By intersecting multi-omics signals, CARDIOWAS effectively narrows down large gene lists into actionable targets, accelerating translational insights in cardiovascular disease.

**Figure 3.**
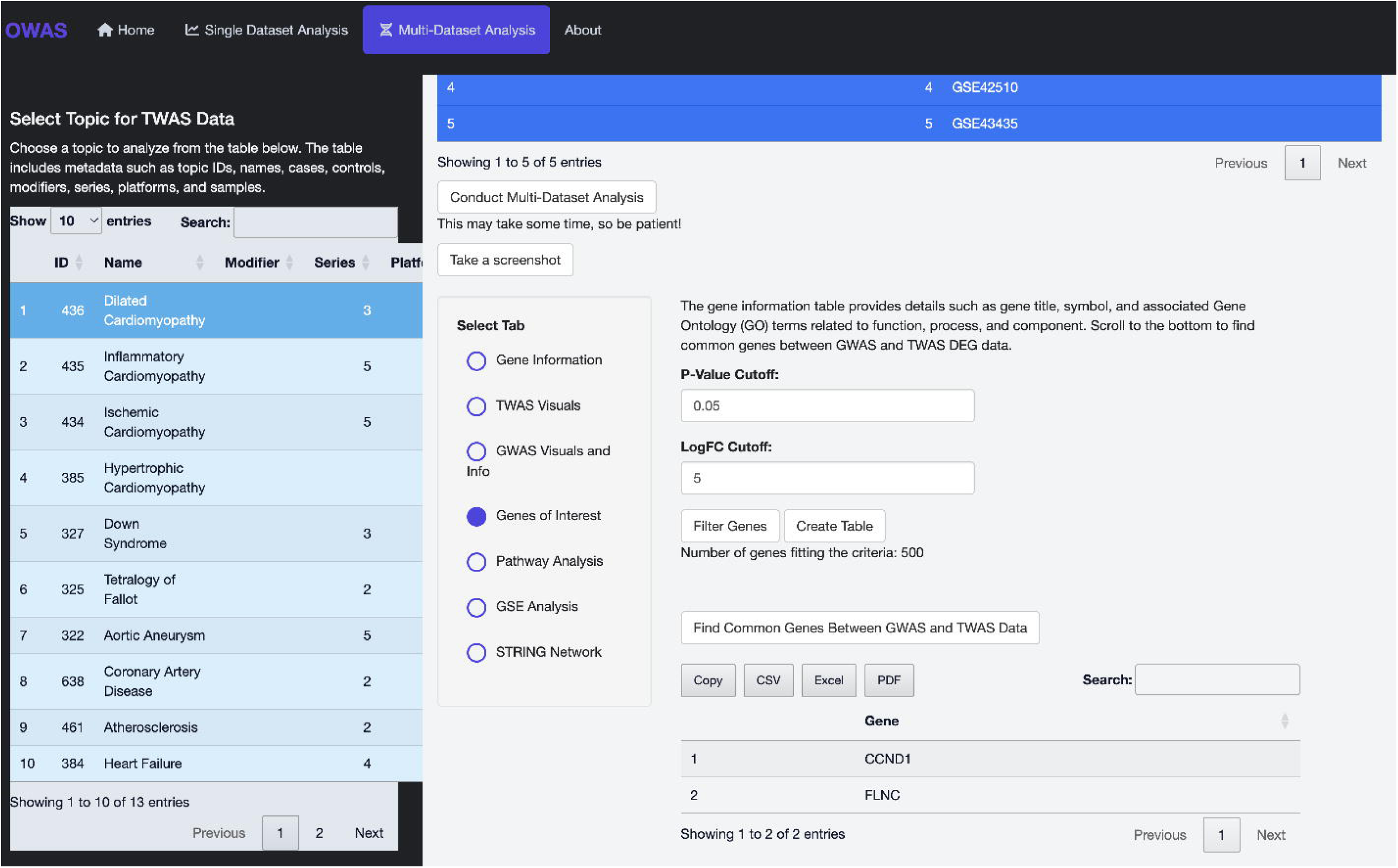
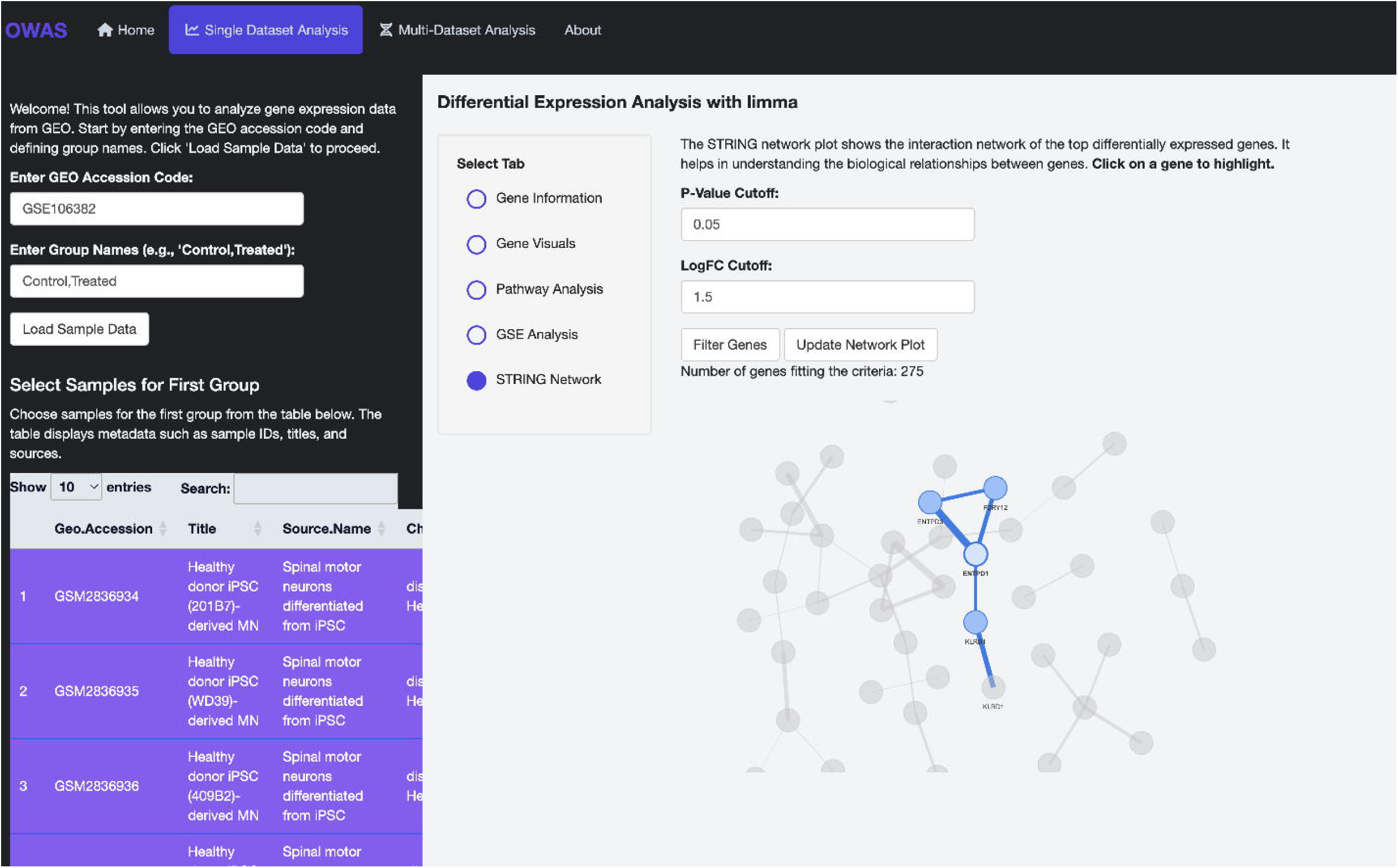
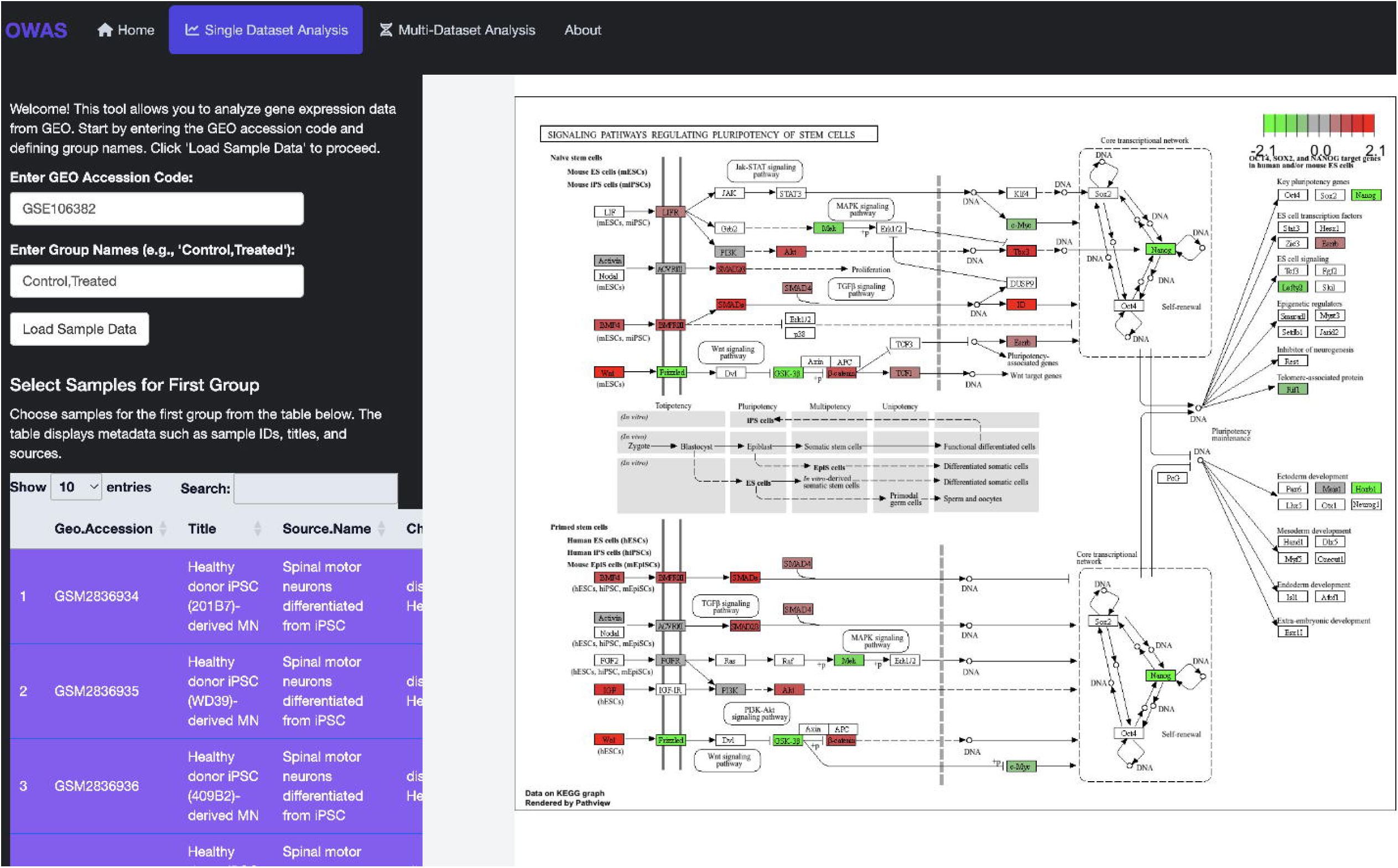
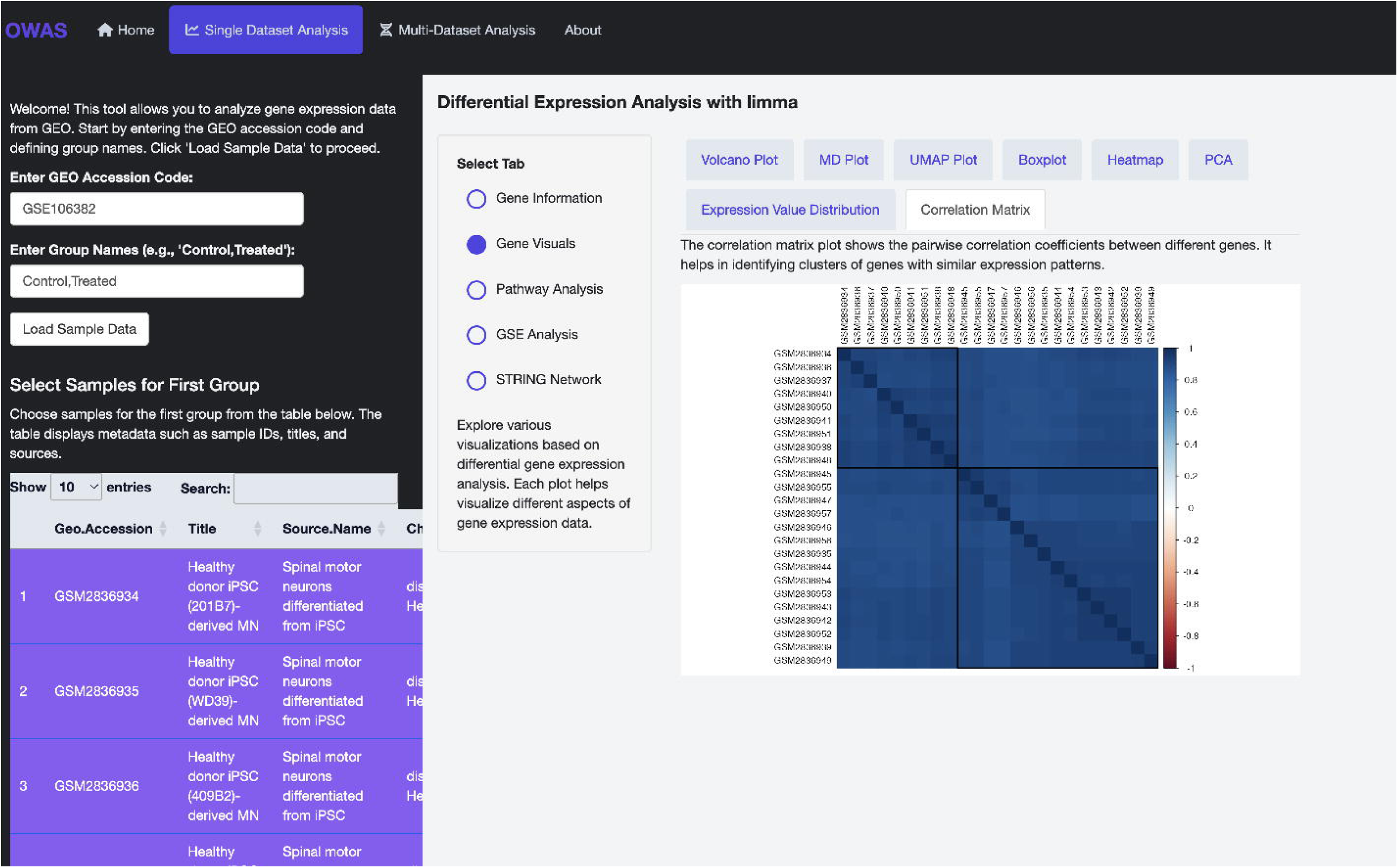
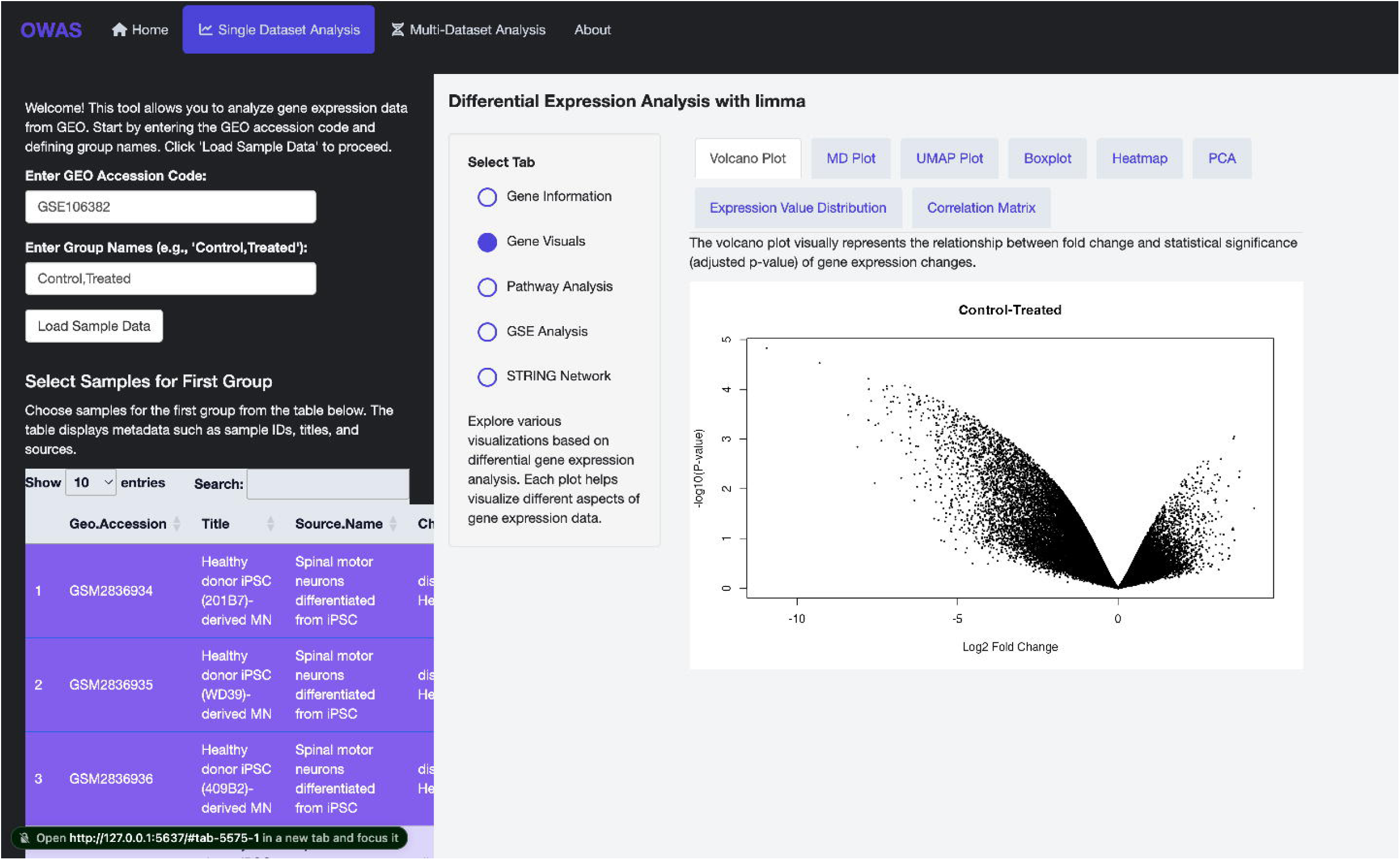
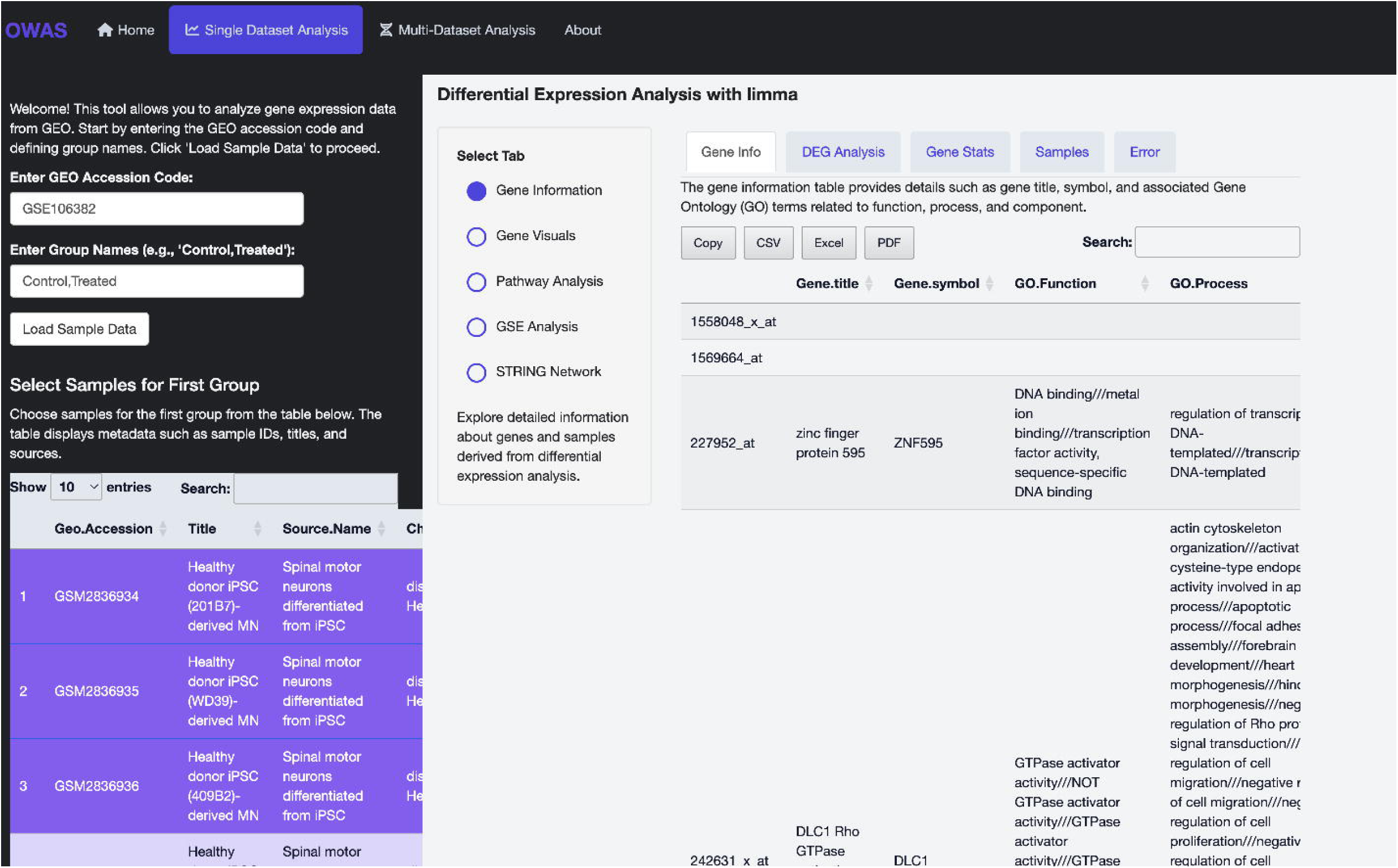
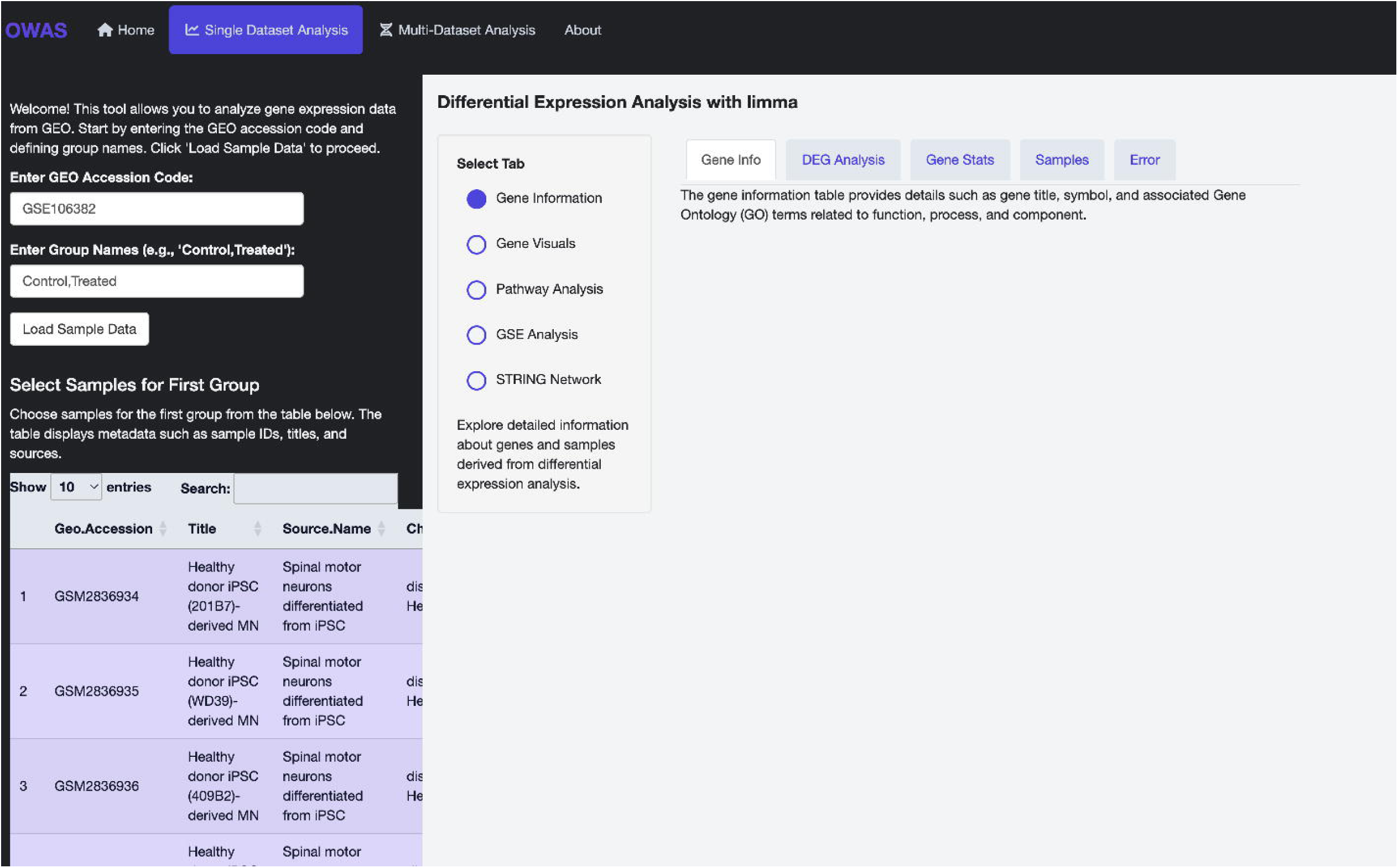
Multi-dataset integration and GWAS overlay in CARDIOWAS for cardiovascular disease analysis. (a) Interface for selecting disease topics (e.g., dilated cardiomyopathy) to perform TWAS-based multi-dataset analysis. Users can browse available conditions and view associated metadata including sample size, series, and platforms. (b) Dataset selection screen showing five GEO series associated with dilated cardiomyopathy. These datasets are harmonized for batch correction and meta-analysis upon launching the analysis. (c) Heatmap of the top 20 differentially expressed genes across all integrated datasets. Color intensity represents normalized expression levels, highlighting gene clusters and sample relationships. (d) Manhattan plot displaying genome-wide SNP-trait associations for dilated cardiomyopathy. Peaks correspond to loci with significant associations across chromosomes, aiding in the identification of candidate genomic regions. (e) Pie chart illustrating the proportion of SNPs falling into functional categories such as intergenic, intronic, missense, and regulatory region variants. This provides biological context for the variant landscape. (f) Annotated Manhattan and region plot highlighting user-specified genes (*IL23R, NOTCH4, NOD2*) and zooming into the *PRKCA* locus. This enables focused interpretation of functional regions with high association signals. (g) Table of overlapping genes identified between TWAS and GWAS analyses, filtered by user-defined thresholds. In this example, *CCND1* and *FLNC* were found to be shared across expression and variant-based analyses, representing high-priority targets for downstream investigation.

## Discussion

The development of CARDIOWAS (Omics-Wide Association Analysis Platform) marks a significant advancement in the field of cardiovascular systems biology by bridging transcriptomic and genomic datasets within an interactive, user-friendly environment. Traditional approaches to RNA-seq analysis often focus on identifying differentially expressed genes (DEGs) without accounting for the broader context of pathway-level changes or the potential contribution of genetic variants to transcriptional dysregulation. CARDIOWAS addresses this limitation by providing integrated modules for DEG detection, pathway enrichment, and multi-cohort analysis, as well as overlaying transcriptomic data with genome-wide association study (GWAS) results. In our analysis of single datasets, CARDIOWAS allowed detailed visualization and interpretation of individual study results. The incorporation of volcano plots, gene ontology tables, correlation matrices, STRING interaction networks^19^, and KEGG pathway enrichment tools empowers users to derive both gene-level and systems-level insights. For example, in dataset GSE106382^20^, we identified distinct transcriptomic signatures linked to immune regulation and vascular remodeling, both key contributors to heart failure and coronary artery disease (CAD) pathophysiology. The ability to conduct this level of analysis without requiring programming expertise represents a major step toward democratizing access to omics analytics in cardiovascular research. CARDIOWAS’s multi-dataset analysis functionality extends its utility beyond single-cohort studies by harmonizing multiple datasets under a unifying disease phenotype. Using dilated cardiomyopathy as a use case, CARDIOWAS successfully integrated five GEO datasets spanning different platforms and sample sources. The resulting meta-analysis produced highly robust DEG profiles, as evidenced by the consistent gene expression patterns captured in heatmaps and correlation matrices. This approach enhances statistical power and reduces dataset-specific noise, providing more reliable candidate genes and pathways for hypothesis generation. Perhaps one of the most innovative features of CARDIOWAS is its seamless integration of GWAS data. The platform allows users to explore the genomic architecture of disease by visualizing Manhattan plots, identifying functional variant classes, and narrowing candidate regions of interest through interactive region plots. In our example, the integration of TWAS and GWAS results for dilated cardiomyopathy identified overlapping genes such as *CCND1*^21^ and *FLNC*^22^, which are supported by both transcriptomic dysregulation and SNP association. This highlights the power of CARDIOWAS to uncover convergent evidence across omics layers, a critical step toward precision medicine applications. Additionally, CARDIOWAS’s support for user-defined SNP annotation thresholds and gene highlighting allows researchers to tailor analyses to specific hypotheses, such as identifying novel regulatory variants or validating known pathogenic loci. The SNP-type classification module also provides a deeper understanding of the molecular mechanisms driving disease by distinguishing between structural, regulatory, and non-coding variation. Despite these strengths, some limitations remain. Currently, somatic mutation detection is not yet integrated into the platform. This capability is essential for studying clonal hematopoiesis and mosaicism, which are increasingly recognized as important contributors to cardiovascular disease, especially in aging populations. Additionally, while CARDIOWAS currently supports TWAS and GWAS integration, future iterations could incorporate proteomic and epigenomic data to further enhance pathway resolution and capture multilevel biological interactions. Looking forward, CARDIOWAS will continue to evolve as a translational research tool. Planned upgrades include the integration of somatic variant identification from RNA-seq data, drug-target prediction based on pathway perturbation, and patient-specific model testing through custom data upload. These enhancements will empower researchers and clinicians to use CARDIOWAS not only for discovery science but also for individualized risk stratification and therapeutic guidance. In conclusion, CARDIOWAS represents a powerful and accessible platform for multi-omics integration in cardiovascular disease research. Its ability to link transcriptomic alterations with genomic architecture at the pathway level offers unprecedented potential for understanding disease mechanisms, identifying biomarkers, and advancing precision cardiovascular medicine.

### Platform Development

To make the analysis accessible to a broad range of researchers, we developed an interactive Shiny application that enables users to access NCBI GEO datasets and perform custom differential expression analyses, explore pathway-level expression changes, integrate GWAS data to examine transcriptome-variant-pathway associations, and visualize risk stratification and potential drug targets. The interface is designed for intuitive use with minimal user intervention, allowing both computational and experimental researchers to uncover biologically meaningful insights with ease.

### Future Development

The next iteration of CARDIOWAS is designed to further enhance its translational potential. Planned modules include the identification of somatic mutations directly from RNA-seq data, pathway enrichment analyses driven by somatic variants, prediction of drug-target relationships based on transcriptomic perturbations, and the ability to test patient-specific models using uploaded transcriptome profiles. These upgrades will support comprehensive omics-driven stratification and therapeutic planning for cardiovascular diseases.

## Conclusion

CARDIOWAS represents a significant advancement in integrative omics analysis for cardiovascular research. By combining transcriptomics and genomics at the pathway level, the platform offers unique insights into disease mechanisms, risk stratification, and personalized therapeutic planning. CARDIOWAS is positioned to become a powerful tool in translational cardiovascular research and precision medicine.

### Availability

The CARDIOWAS Shiny app is accessible via https://github.com/krishalok/CardiCARDIOWAS

## Acknowledgements

We acknowledge the contributors of the NCBI GEO and GWAS Catalog datasets, and the patients who provided samples for these studies.

